# Overall patterns of eye-specific retino-geniculo-cortical projections to layers III, IV and VI in primary visual cortex of the prosimian galago, and correlation with cytochrome oxidase blobs

**DOI:** 10.1101/2022.06.08.495376

**Authors:** Jaime F. Olavarria, Huixin Qi, Toru Takahata, Jon H. Kaas

## Abstract

Studies in galago have not provided a comprehensive description of the organization of eye specific retino-geniculate-cortical projections to the recipient layers in V1. Here we demonstrate the overall patterns of ocular dominance domains in layers III, IV and VI revealed after injecting the transneuronal tracer wheat germ agglutinin conjugated to horseradish peroxidase (WGA-HRP) into one eye. We also correlate these patterns with the array of cytochrome oxidase (CO) blobs in tangential sections through the unfolded and flattened cortex. In layer IV, we observed for the first time that eye-specific domains form an interconnected pattern of bands 200-250 um wide arranged such that they do not show orientation bias and do not meet the V1 border at right angles, as is the case in macaques. We also observed distinct patterns of ocular dominance patches in layer III and layer VI. The patches in layer III, likely corresponding to patches of K LGN input described previously, align with layer IV ocular dominance columns (ODCs) of the same eye dominance. Moreover, the layer III patches overlap partially with virtually all CO blobs in both hemispheres, implying that CO blobs receive K LGN input from both eyes. Finally, we found that CO blobs straddle the border between neighboring layer IV ODCs. These results, together with studies showing that a high percentage of cells in CO blobs are monocular, suggest that CO blobs consist of ipsilateral and contralateral subregions that are in register with underlying layer IV ocular dominance columns of the same eye dominance. In macaques and humans, CO blobs are centered on ODCs in layer IV. Our finding that CO blobs in galago straddle the border of neighboring layer IV ODCs suggests that this may represent an alternative way by which visual information is processed by eye specific modular architecture in mammalian V1.

## Introduction

In the prosimian galago (bush baby), as in other primates, retinal input is relayed to the lateral geniculate nucleus (LGN) and primary visual cortex (V1) along three major parallel pathways named after the distinct class of retinal ganglion cells from which they originate. These pathways are the magnocellular (M), parvocellular (P), and koniocellular (K) pathways (reviewed in Casagrande & Kaas, 1994). Studies of the projections of these three pathways in galago V1 revealed that projections of LGN M and P cells terminate with the upper and lower divisions of layer IV, respectively, with a minor projection to layer VI, whereas K LGN cells project to layers III and I (Lachica & Casagrande, 1992; Casagrande & Kaas, 1994, Ding & Casagrande, 1997).

Previous anatomical studies in galago provided preliminary evidence of periodicity in layer IV (Casagrande & DeBruyn, 1982; Diamond et al., 1985; Glendenning et al., 1976), and physiological studies suggested the existence of ocular dominance columns (ODCs) in galago based on the observation that cells with similar ocular dominance responses were grouped together (Debruyn et al., 1993). More recently, an optical imaging study in galago revealed clusters of monocular responses in galago (Xu et al., 2005), but because this imaging method cannot discern patterns of activity from separate layers across the depth of the cortex, the results do not inform about the patterns of LGN projections to specific layers.

It is well established that K LGN cells project in a patchy fashion to layer III in galago V1 (Casagrande & DeBruyn, 1982, Diamond et al., 1985, Lachica & Casagrande, 1992), but at present it is not known whether the overall patterns of LGN projections to layers IV and VI form separated patches as in layer III, or stripe-like patterns resembling the patterns of ODCs in macaque monkey (Tootell et al., 1988; Florence & Kaas, 1992; Horton & Hocking, 1996b; Olavarria & Van Essen, 1997), or interconnected bands as in other primate species (Takahata et al., 2014) and cats (Anderson et al., 1988). Similarly, it is not known whether layer III patches are randomly distributed across the expanse of V1, or instead are aligned with layer IV domains of the same eye dominance. This question is of interest given that direct projections to layers III and IV originate from different LGN cell types. Finally, while it has been reported that single axon projections from K LGN cells arborize within cytochrome oxidase (CO) blobs in galago V1 (Lachica & Casagrande, 1992), the relationship between CO blobs and ODCs in galago layer IV has not been examined with anatomical methods. Optical imaging studies in galago (Xu et al., 2005), owl monkey (Kaskan et al., 2007) and New World marmoset (Roe at al., 2005) found no clear relationship between ocular dominance domains and CO blobs, whereas studies in macaque monkey have established that CO blobs are consistently centered on ODCs (Horton, 1984; Blasdel & Salama, 1986; Tootell et al., 1988; Bartfeld & Grinvald, 1992, Yoshioka et al., 1996). Here we address these standing questions in the prosimian galago with the ultimate goal of advancing our understanding of the organization and evolution of the visual system in primates, including humans. To this end, we injected the transneuronal tracer wheat germ agglutinin conjugated with horseradish peroxidase (WGA-HRP) into one eye and examined and correlated the patterns of WGA-HRP and CO labeling in tangential histological sections through the unfolded and flattened cortex.

In layer III, we observed that the WGA-HRP labeling appears as an array of distinct patches that most likely represent the patchy K input to layer III shown previously in galago (Lachica & Casagrande, 1992) and owl monkey (Ding & Casagrande, 1997). In layer VI, we observed distinct patches that appear somewhat larger than the patches in layer III, but of similar density. In granular cortex, the labeling adopts the form of interconnected bands or stripes whose overall array is reminiscent of the pattern of ODCs in others primates (e.g., owl monkey [Takahata et al., 2014]) and in cats (Anderson et al., 1988). Moreover, we found that the patches in layer III are aligned with ODCs in layer IV of the same eye dominance, and that virtually *all* CO blobs overlap with WGA-HRP labeled patches in layers III. Since only one eye was injected with WGA-HRP, these data imply that all CO blobs receive K LGN input from both eyes, and suggest that these inputs remain largely segregated within CO blobs, consistent with previous studies in galago indicating that a high percentage of cells recorded within CO blobs are monocular (DeBruyn et al., 1993). Moreover, we found that CO blobs typically span the borders between left and right eye columns. Based on these findings, we propose a novel model in which CO blobs straddle the border between neighboring ipsilateral and contralateral ODCs in layer IV, such that the resulting CO sub-regions receive segregated K LGN input of the same eye dominance as the underlying ODCs.

## Materials and Methods

### Animals

The data on the patterns of ocular domains and CO blobs in the present study come from a total of 7 hemispheres from 4 adult galagos (Galago crassicaudatus) weighing 800–1300 g. All procedures were performed in the Psychology Department of Vanderbilt University, following protocols approved by the Animal Care and Use Committee at Vanderbilt University. The research was conducted in accordance with the animal care guidelines of the National Institutes of Health.

### Intravitreal injections

Galagos received monocular intravitreal injections containing 1.5 mg of WGA-HRP (Sigma-Aldrich, St Louis, MO) dissolved in 20 µL of normal saline. The tracer was delivered during 15- 20 min through glass micropipettes driven to about 5-6 mm deep into the posterior chamber of one eye. Anesthesia was induced with Ketaset (0.05 mg/Kg, im) and maintained with isoflurane 2%. Arterial pulse oxygenation, pulse rate, respiration rate, and core body temperature were monitored throughout the surgical procedures.

### Tissue processing

After a survival period of 4-5 days, the animals were deeply anesthetized with pentobarbital sodium (100 mg/kg i.p.) and perfused through the left cardiac ventricle with 0.9% saline until the fluid from the right atria was clear, followed by a brief (7-8 min) perfusion with 2% paraformaldehyde in 0.1 M phosphate buffer (pH 7.4). The brains were removed and photographed. Visual cortex was unfolded, flattened (Olavarria & Van Sluyters, 1985) and cut in the tangential plane (50 µm in thickness) in a freezing microtome.

### Histological procedures

The first 4-5 cortical tangential sections through visual cortex were reacted for CO histochemistry according to the protocol (Wong-Riley, 1979). The remaining tangential sections were reacted for HRP histochemistry with tetramethyl benzidine as the chromogen (Mesulam, 1978).

### Data acquisition and analysis

Individual sections containing CO or WGA-HRP labeling were scanned (Epson 4990), and the resulting digital images were used to reconstruct the patterns of CO and HRP labeling in the primary visual cortex. The data were analyzed using Photoshop CS5 (Adobe Systems, San Jose, CA), and all image processing used, including contrast enhancement and intensity level adjustments, were applied to the entire images and never locally. Patterns of ODCs or CO over large areas were reconstructed by superimposing two or more sections in Photoshop CS5 (Adobe Systems, Inc. CA). Images in separate Photoshop layers were carefully aligned with each other by using the edges of the sections, blood vessels, and other fiducial marks, and the final pattern of ODCs or CO were obtained by merging all images together. To facilitate obtaining thresholded images of the final ODC and CO patterns, images were high-pass filtered to remove gradual changes in staining density. To correlate the patterns of CO and ODCs, these were carefully aligned in Photoshop using radial blood vessels, tissue borders and penetrating marks made in the flattened cortex in regions close to V1. To calculate the density and size of CO blobs, labeled patches in layers III and VI, thresholded images of these elements were analyzed using ImageJ. To calculate the density of a certain class of elements, we counted the number of elements in the sampled region, and then divided the number obtained by the total area of the sampled region. To measure the average size of a certain class of elements, we measured the total area occupied by a number of elements in each element class, and divided it by the number of elements in the sampled area (Table 1). To test whether labeled patches in layer III are aligned with ODCs of the same eye dominance in layer 4, we performed a χ^2^ test, df = 1, significance set at P< 0.05.

**Table 1.**
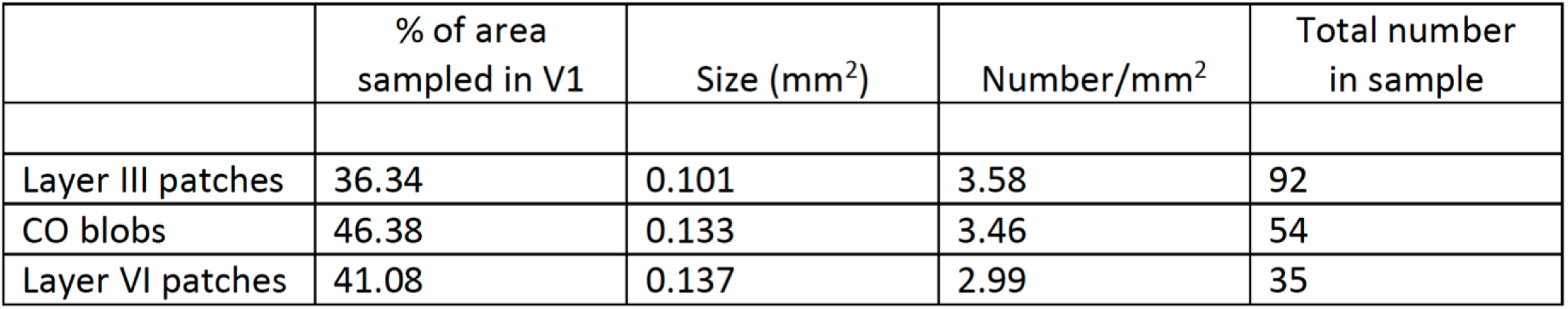
Coverage, size and density of layer Ill patches, CO blobs and layer VI patches

|  | % of area<br>sampled in V1 | Size (mm<sup>2</sup>) | Number/mm<sup>2</sup> | Total number<br>in sample |
| --- | --- | --- | --- | --- |
| Layer III patches | 36.34 | 0.101 | 3.58 | 92 |
| CO blobs | 46.38 | 0.133 | 3.46 | 54 |
| Layer VI patches | 41.08 | 0.137 | 2.99 | 35 |

## Results

To investigate the distribution of retino-geniculo-cortical input to striate cortex (V1) in galagos, we injected the transneuronal tracer WGA-HRP into one eye and analyzed the patterns of WGA labeling in tangential sections trough supragranular, granular and infragranular layers of V1 in both hemispheres. The transneuronal anterogradely transport properties of WGA-HRP have been demonstrated at both the optical and structural level (Itaya & van Hoesen, 1982, Itaya et al., 1984). These properties ensure that the cortical labeling in these cortical targets corresponds to direct LGN projections (Iataya et al., 1984). In addition, we correlated the WGA-HRP labeling patterns with the distribution of CO blobs revealed in a separate set of tangential sections trough supragranular cortex (Condo and Casagrande, 1990). We identified the cortical layers displaying WGA-HRP labeling by comparing our results with previous descriptions of retino-geniculate-cortical projections in galago revealed in parasagittal or coronal histological sections (Casagrande and DeBruyn, 1982, Gledenning et al., 1976), or in tangential sections through V1 (Lachica and Casagrande, 1992). We describe the projection patterns in supragranular layer (layer III), granular layer (layer IV) and infragranular layer (layer VI), the correlation between WGA-HRP labeling patterns in layers III and IV, and the correlation of CO blobs with labeling patterns in layers III and IV.

The location of the area of galago V1 that is visible on the surface of the occipital lobe is indicated by the dotted line in Fig. 1A. V1 extends ventrally over the tentorium and medially on the dorsal and ventral banks of the calcarine fissure (Rosa et al., 1997). Unfolding and flattening the entire V1 required cutting along the fundus of the calcarine fissure (Fig. 1B). The rectangle in Fig. 1B indicates the region of V1 illustrated in Figs. 1C, D, E, and the box inside the rectangle outlines the common region analyzed in these images. Figure 1C,D shows the labeling pattern in layers III and IV, respectively, in V1 ipsilateral to the WGA-HRP eye injection in galago 7, and Fig. 1E shows the patterns of CO blobs in the same region. For each pattern, a thresholded version of the common region is shown in color below each image (Fig. 1F,G,H), and the relationships between these patterns in this case are illustrated in Figure 2.

**Figure 1.**
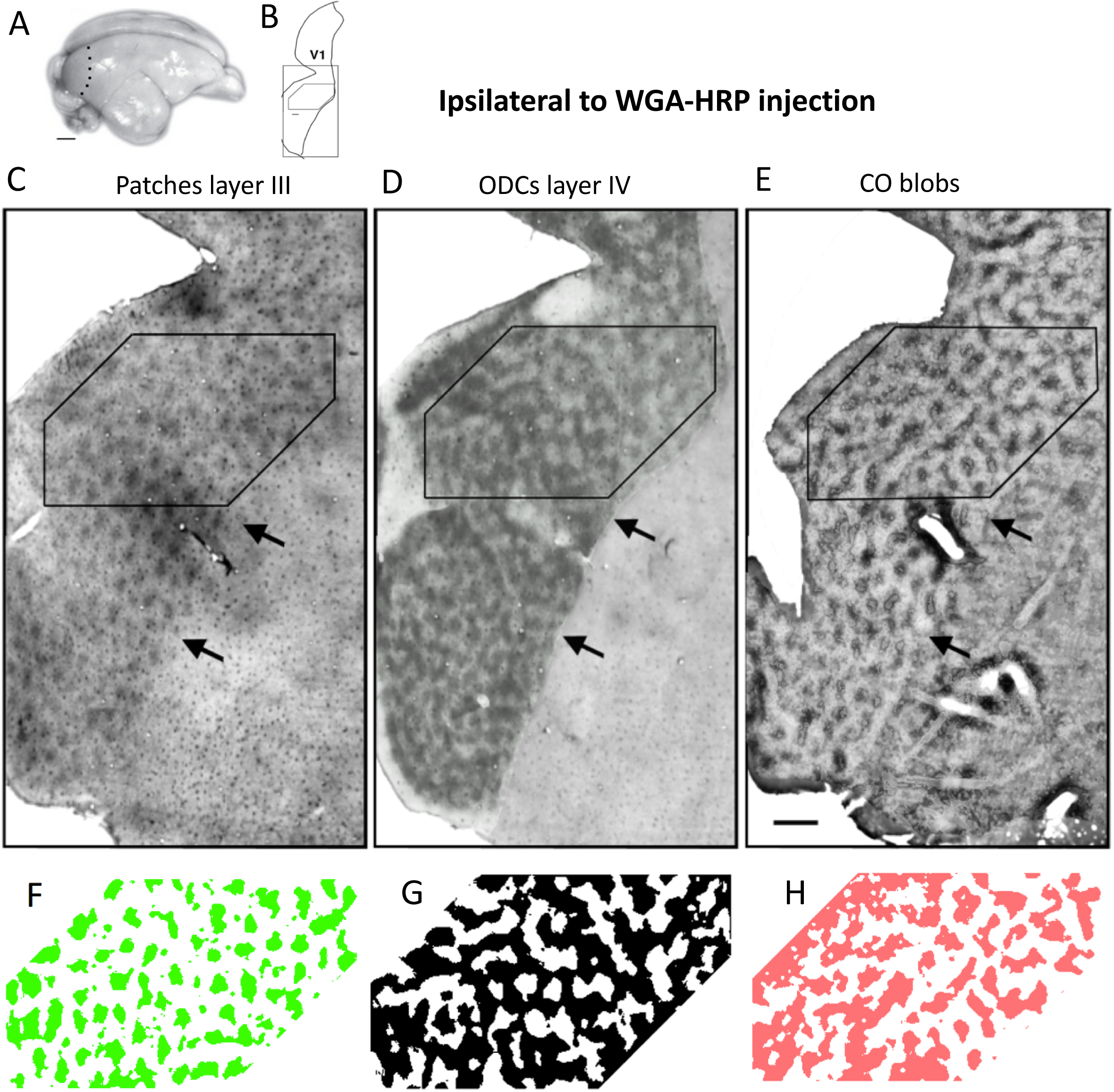
WGA-HRP labeling in layers III and IV ipsilateral to the intraocular injection of WGA-HRP, and CO blobs in supragranular layers in galago 7. A. Galago brain showing the location of V1 in the occipital cortex of the right hemisphere. Dotted line indicates anterior border of V1. B. The rectangle indicates the region of the unfolded and flattened V1 illustrated in Figs. 1C, D, E, and the box inside the rectangle outlines the common region analyzed in these images. C,D. Labeling pattern in layers III and IV, respectively. E. Pattern of CO blobs in the same hemisphere. Arrows in C,D,E indicate anterior border of V1. F,G,H. Thresholded colored versions of the patterns in C,D,E, respectively. Scale bar in E = 1.0 mm.

**Figure 2.**
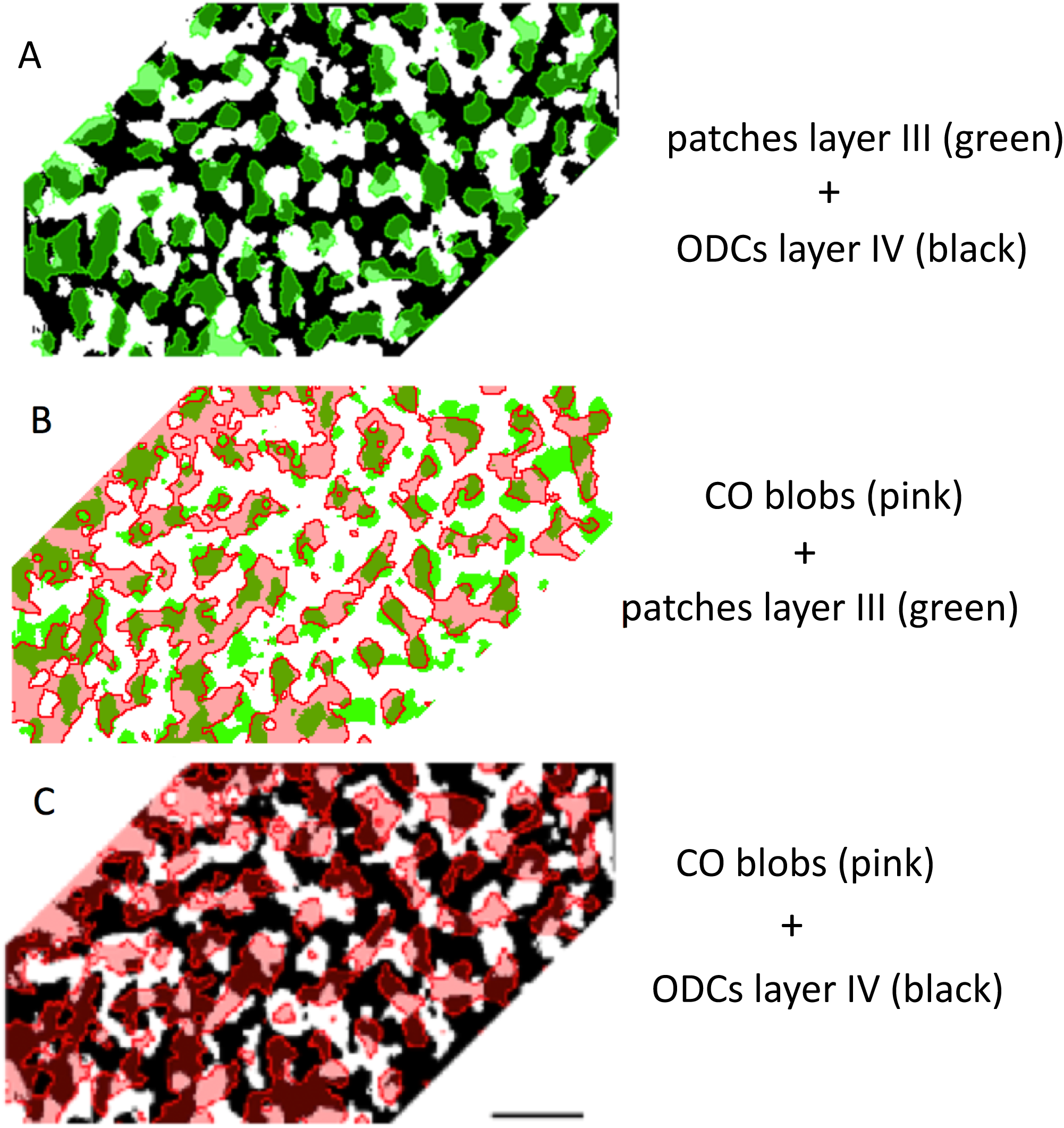
Relationships between the WGA-HRP and CO labeling patterns in V1 ipsilateral to the eye injection in galago 7 shown in Fig 1F,G,H. A. Superimposition of the labeling pattern in layer III (green patches) over the pattern of ODCs in layer IV (black). B. Superimposition of the pattern of CO blobs (pink) over the labeling pattern in layer III (green). C. Superimposition of the pattern of CO blobs (pink) over the labeling of ODCs in layer IV (black). Scale bar = 1.0 mm.

### Retino-geniculo-cortical projections to layer III in V1

Projections to layer III were examined in hemispheres ipsilateral and contralateral to the WGA-HRP eye injection. In both hemispheres, WGA-HRP labeled projections to layer III appear as arrays of distinct patches. Figures 1C and 3A illustrate ipsilateral layer III projections in galago 7 and galago 5, respectively. Patchy contralateral projections to layer III are illustrated in the section in Fig. 5A,B (galago 5), as well as in the immediately deeper section from the same case (Fig. 6A). Most of Fig. 6A illustrates the patchy labeling pattern in layer III, except in the ventral region (white rectangle), which shows labeling in layer IV (described below). In this ventral region, it is interesting to notice that in both Figs. 5A and 6A, the labeling in the area marked with an asterisk is uniformly dense. This region represents the peripheral monocular segment innervated by the contralateral eye, as described in other primates (Horton and Hocking, 1966b), and the arrow next to the asterisk in Figs. 5A and 6A indicates the border of peripheral V1.

**Figure 3.**
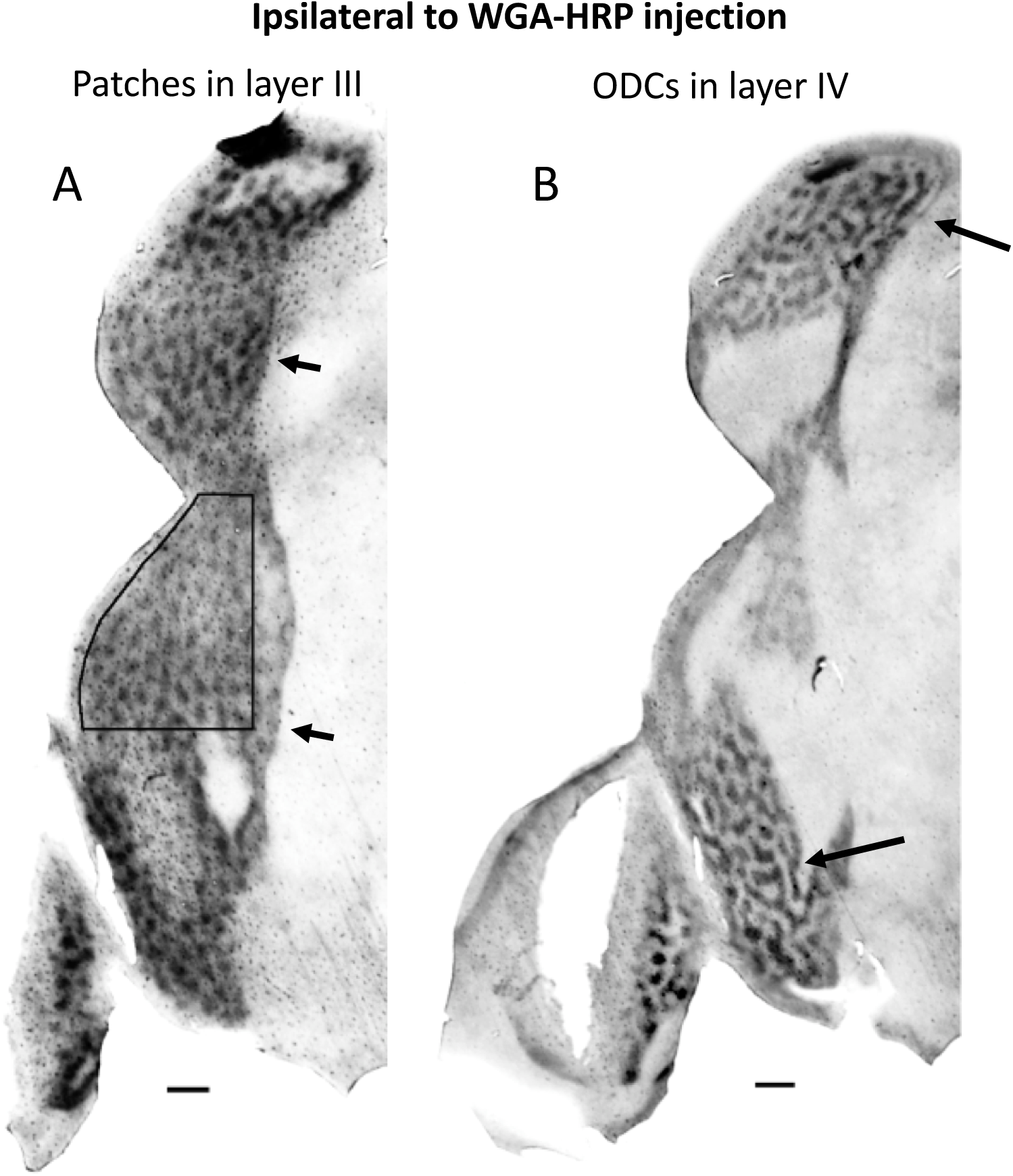
Patterns of WGA-HRP labeling in layers III and IV in the hemisphere ipsilateral to the eye injection of WGA-HRP in galago 6. A. Labeling in layer III. Arrows indicate the anterior border of V1. The box indicates region analyzed in Fig. 4. B. Labeling pattern in layer IV in regions indicated by the arrows. Scale bar = 1.0 mm.

The labeled patches in layer III occupy 36.34% of the area analyzed in central V1. On average, they measure 0.101 mm2 in area size, and the number of patches per unit area is 3.58 patches/mm2. The patchy labeling in layer III most likely represent the patchy koniocellular input to layer III shown previously (Diamond et al., 1985; Lachica & Casagrande, 1992). Patchy labeling was not observed beyond the anterior border of V1, indicated by arrows in Figs. 1C, 3A.

### Retino-geniculo-cortical projections to layer IV in V1

The distribution of projections to layer IV is strikingly different from that to layer III. Instead of isolated patches, the labeling in layer IV in both hemispheres forms patterns of interconnected bands or stripes 200-250 um in width. Figure Fig. 1D illustrates the labeling pattern in layer IV in the hemisphere ipsilateral to the eye injection in galago 7. A similar pattern in layer IV was observed in the hemisphere ipsilateral to the eye injection in galago 5 (long arrows in Fig. 3B), as well as in the hemisphere contralateral to the eye injection (Fig. 6B). Figure 6B shows a magnified view of the area outlined by the white rectangle in Fig. 6A.

The pattern of ocular dominance columns (ODCs) we observed in galago layer IV resembles the pattern of ODCs in the cat layer IV (Anderson et al., 1988), squirrel monkey (Horton and Hocking, 1996a) and owl monkey (Takahata et al., 2014). Moreover, as in these species, the bands or stripes in galago layer IV do not appear to have a preferred orientation across V1, and, in particular, they do not tend to meet the border of V1 at right angles, as in macaque monkeys (Florence and Kaas,1992; Horton and Hocking, 1996b; Fig. 6 in Olavarria and Van Essen, 1997). Ipsilateral labeled layer IV ODCs occupy 60,59% of the area examined, while contralateral labeled layer IV ODCs occupy 58.88% of the area examined, suggesting some overlap between right and left ODCs in layer IV.

### Relation between layer III patches and layer IV ODCs

Layer III patches in galago have been revealed after tracer injections into eye specific K LGN layers (Lachica and Casagrande, 1992), following transneuronal transport of tracers after monocular tracer injections (Casagrande and Debruyn, 1982), and patchy activation patterns have been revealed with imaging techniques following monocular stimulation in galago (Xu et al., 2005) and owl monkey (Kaskan et al., 2007). A question not addressed by previous studies is whether layer III patches are randomly distributed in V1, or whether they are aligned with layer IV bands of the same eye dominance. Figure 2A shows the superimposition of the pattern of layer III patches over the pattern of ODCs in layer 4 from the case in Fig. 1. Inspection of Fig. 2A suggests that layer III patches (green) are preferentially aligned with layer 4 ODCs (black). To address this issue quantitatively, we performed a χ^2^ analysis of the distribution of labeling in layers III and IV in both hemispheres following an injection of WGA-HRP into one eye of galago 7. For a random distribution of layer III patches, the proportion of labeled layer III patches over labeled and unlabeled ODCs in layer IV should be equal to the proportion of areas covered by labeled and unlabeled ODCs, respectively. Our analysis showed that the proportion of ipsilateral labeled layer III patches over ipsilateral labeled layer 4 ODCs was greater than expected (74.58%; expected 60.59%), while the proportion of ipsilateral labeled layer III patches over unlabeled contralateral layer 4 ODCs was smaller than expected (25.42%; expected 39.40%), χ^2^ = 8.19, df = 1, P < 0.01. Similarly, the proportion of contralateral labeled layer III patches over contralateral labeled layer 4 ODC was greater than expected (70.758%; expected 58.887%), while the proportion of contralateral labeled layer III patches over unlabeled ipsilateral layer IV ODCs was smaller than expected (29.241%; expected 41.112%), χ^2^ = 5.82, df = 1, P < 0.05. These results reject the null hypothesis that labeled layer III patches are distributed randomly over V1, and suggest that the radial registration of eye specific geniculo-cortical projections to different cortical layers is largely maintained even when the direct projections to these layers originate from different cells types in the LGN.

### Retino-geniculo-cortical projections to layer VI in V1

It is known that in galagos, as in macaques, P and M geniculate cells send relatively minor projections to layer VI (reviewed in Casagrande and Kaas, 1994), but the overall pattern of these projections is not currently known. In the New World spider monkey, Florence et al. (1986) studied ocular segregation in striate cortex following intraocular injections of tritiated proline in one eye. They found that the projections to layer VI are considerably lighter than those to layer IV and do not show clear evidence of ocular periodicity. We found that retino-geniculate projections to galago layer VI form patches that are somewhat less distinct than the patches in layer III. Figure 7A,B shows labeled patches in layer VI ipsilateral and contralateral to the WGA-HRP injection, respectively, in galago 7, anterior is to the right in both images. The labeled patches in layer VI occupy 41.08% of the area analyzed in central V1, and on average, they measure 0.137 mm2 in area size, and the number of patches per unit area is 2.99 patches/mm2. Compared with labeled patches in layer III, layer VI patches appear to be somewhat larger, cover a larger area of V1, and their number per unit area is smaller (Table 1). It will be interesting to determine whether layer VI patches segregate by eye dominance and whether they correlate with patches in layer III, ODCs in layer IV, and CO blobs in layer VI (Condo and Casagrande, 1990). In the cat, injections of tritiated proline into LGN layers (LeVay and Gilbert, 1976), or WGA-HRP injections into one eye (Olavarria, unpublished results) reveal columnar projection to layer VI that are in register with ODCs in layer IV.

### Correlation between CO blobs and layer III WGA-HRP labeled patches

In macaque and squirrel monkeys, Livingstone and Hubel (1982) reported that injections of either tritiated proline or HRP into the LGN revealed an array of patches that appeared to match the array of cytochrome oxidase blobs in layer III, but it was not resolved whether this projection was direct, or indirect via layer IV. Similar results have been reported in owl monkey (Ding and Casagrande, 1997) and squirrel monkey (Fitzpatrick et al., 1983). In galago, Lachica and Casagrande, 1992) demonstrated that K LGN layers project directly to layer III, and that single axons labeled by injections of WGA-HRP or Phaseolus vulgaris leucoagglutinin (PHA-L) into K LGN layers arborize within CO blobs in layer III (Lachica and Casagrande, 1992). Our results confirm and extend these previous findings in galago by providing evidence that virtually all CO blobs in V1 receive K LGN input from both eyes.

Figure 1E illustrates the distribution of CO blobs in the hemisphere ipsilateral to the WGA-HRP injection in galago 7, and Fig. 2B correlates the distribution of CO blobs (pink) with the distribution of WGA-HRP labeled patches in layer III (green). This figure illustrates that virtually all CO blobs overlap with WGA-HRP labeled patches in layer III. Results from the hemisphere ipsilateral to the eye injection in galago 5 are illustrated in Fig. 4. The pattern of layer III patches (Fig. 4A) and the pattern of CO blobs (Fig. 4B) are superimposed in Fig. 4 (C). Figure 4C shows that virtually all CO blobs (grey) overlap with the patches of K LGN input in layer III (black). Similar results were observed in the hemispheres contralateral to the eye injection. Figure 5A illustrates the distribution of contralateral WGA-HRP labeled patches in layer III in galago 7. The region analyzed is outlined in black and shown at higher magnification in Fig. 5B. Figure 5C shows the pattern of CO blobs in the same region, and Fig. 5D illustrates the correlation between the labeled layer III patches (black) and the pattern of CO blobs (grey) from the same region. Again, we observed that virtually all CO blobs (grey) overlap with layer III patches (black). Moreover, Figs. 2B, 4C and 5D provide evidence that layer III patches typically overlap with only a portion of the CO blobs, and that in some instance they extend beyond the borders of CO blobs, into the interblob region. Together, the results from hemispheres ipsilateral (Fig. 2B, 4C) and contralateral (Fig. 5A) to the eye injection provide strong support to the idea that all CO blobs in galagos receive K LGN input from both eyes. We estimate that CO blobs occupy about 46.38% of the area analyzed in V1. On average, CO blobs measure 0.133 mm2 in size, they occupy 46.38 % if the area sampled in V1, and there are 3.46 blobs/mm2. Comparison of CO blobs with layer III labeled patches suggest that CO blobs occupy a larger area of V1, are larger in size, but the number of CO blobs and layer III patches per unit area is very similar (Table 1).

**Figure 4.**
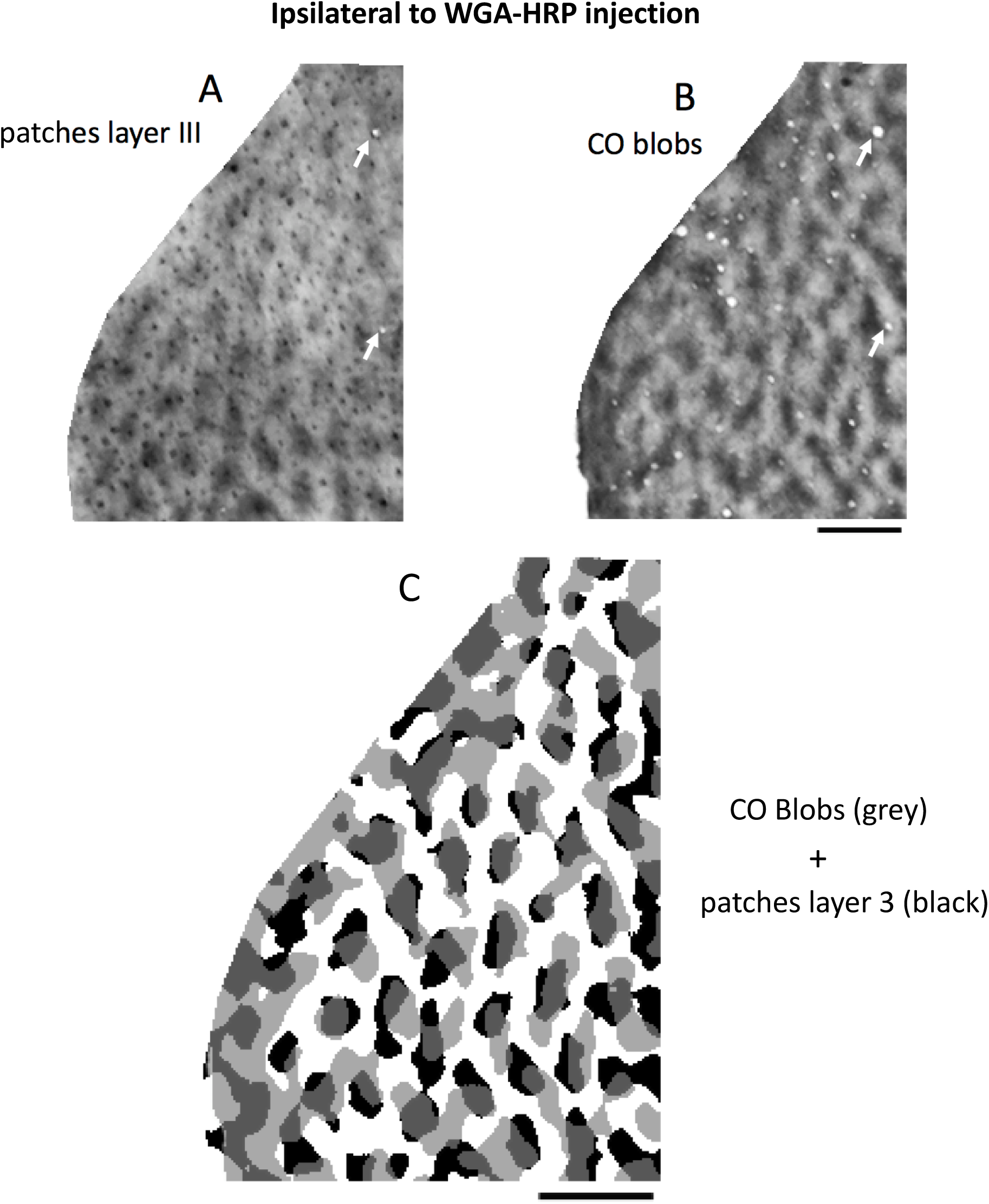
Correlation between the WGA-HRP labeling pattern in layer III and the pattern of CO blobs in hemisphere ipsilateral to the eye injection in galago 6. The analyzed region is indicated by the box in Fig. 3. A. Pattern of labeled patches in layer III. B. Pattern of CO blobs. C. Superimposition of CO blobs (grey) over labeled patches in layer III (black). White arrows in A, B indicate some of the blood vessels used to superimpose patterns in A and B. Scale bars in B,C = 1.0 mm.

**Figure 5.**
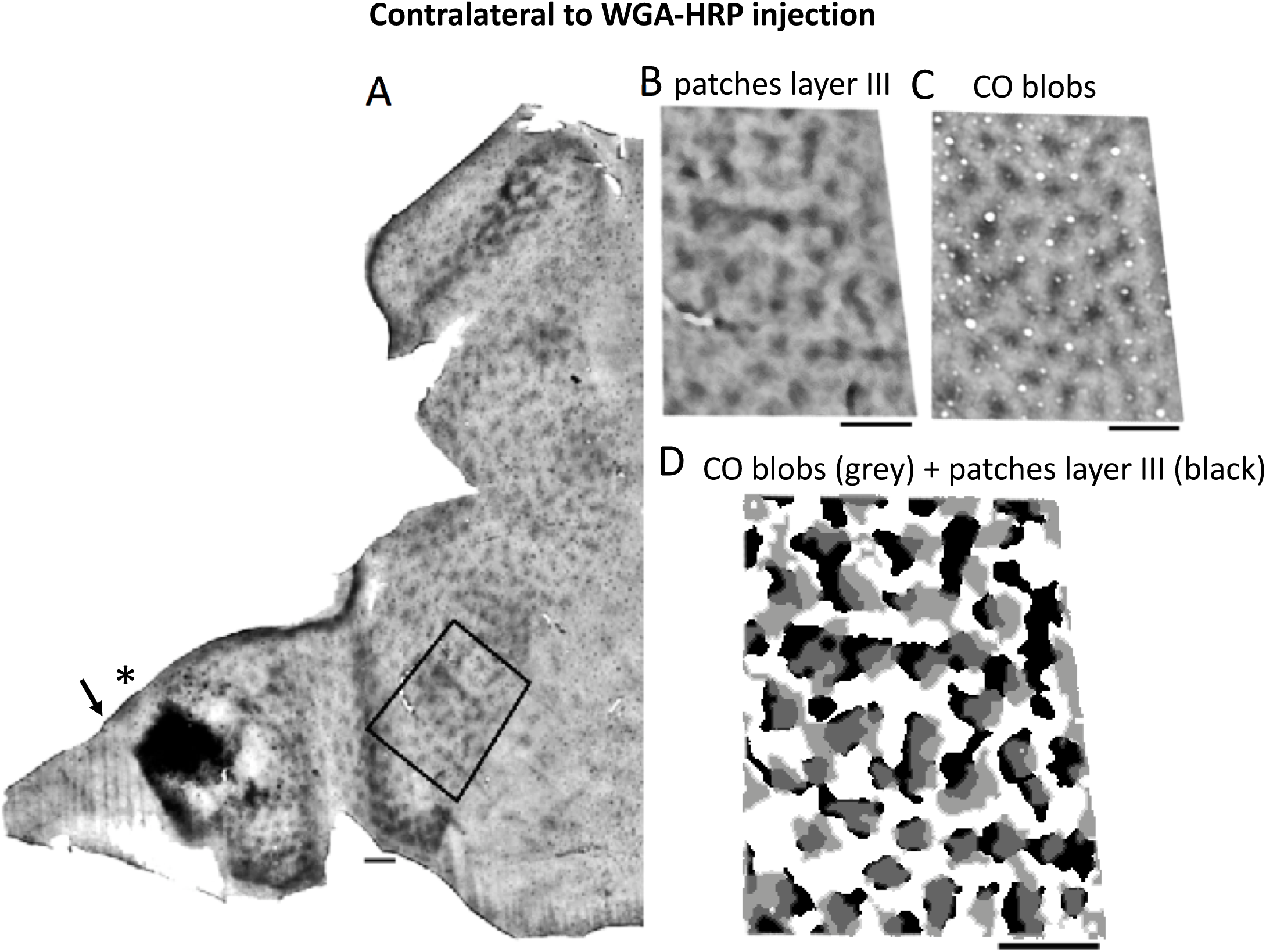
Correlation between the WGA-HRP labeling pattern in layer III and the pattern of CO blobs in hemisphere contralateral to the eye injection in galago 7. A. Overall pattern of labeling in layer III. The analyzed region is indicated by the box in A. B. Pattern of labeled patches in layer III. C. Pattern of CO blobs in the same region. D. Thresholded patterns of CO blobs (grey) over labeled patches in layer III (black) are superimposed. Asterisk in A indicates region of homogeneous labeling in the peripheral monocular segment of V1, and the arrow indicated the peripheral border of V1. Scale bars in A,B,C,D = 1.0 mm

### Correlation between CO blobs and layer IV ODCs

Previous studies have shown that CO blobs are located at the center of ODCs in macaques (Horton, 1984; Blasdel and Salama, 1986; Tootell et al., 1988; Bartfeld and Grinvald, 1992; Yoshioka et al., 1996), but a clear and consistent correlation between CO blobs and ODCs has not been found in New World primates (Roe et al., 2005; Kaskan et al., 2007; Takahata et al., 2014). Similarly, an optical imaging study of intrinsic signals in galago observed no clear relationship between the distribution of CO blobs and the ocular dominance domains (Xu et al., 2005). We found that individual CO blobs consistently straddle the border between neighboring ipsilateral and contralateral layer IV ODCs. Figure 2C (galago 7, ipsilateral to the eye injection) shows the pattern of CO blobs (pink) superimposed on the pattern of ipsilateral (black) and contralateral (white) ODCs, such that virtually each CO blob samples both ipsilateral (black) and contralateral (white) territories. Similar results were obtained in the hemisphere contralateral to the eye injection (Fig. 6, galago 7). The region analyzed is outlined by the white rectangle in Fig. 6A. The labeling pattern in layer IV in this region is enlarged in Fig. 6B, and Figure 6 C shows that pattern of CO blobs in the same region (white arrows indicate common blood vessels). For each of these two patterns, a thresholded version of the common region is shown in black and pink below each image, and the relationship between these patterns is shown in Fig. 6D. As in Figure 2C, Figure 6D shows that virtually all CO blobs (pink) straddle the border between neighboring ODCs, such that each CO blob overlaps with labeled (black) and unlabeled (white) layer IV ODCs. The yellow square outlines the region displayed in Figure 8 (see Discussion).

**Figure 6.**
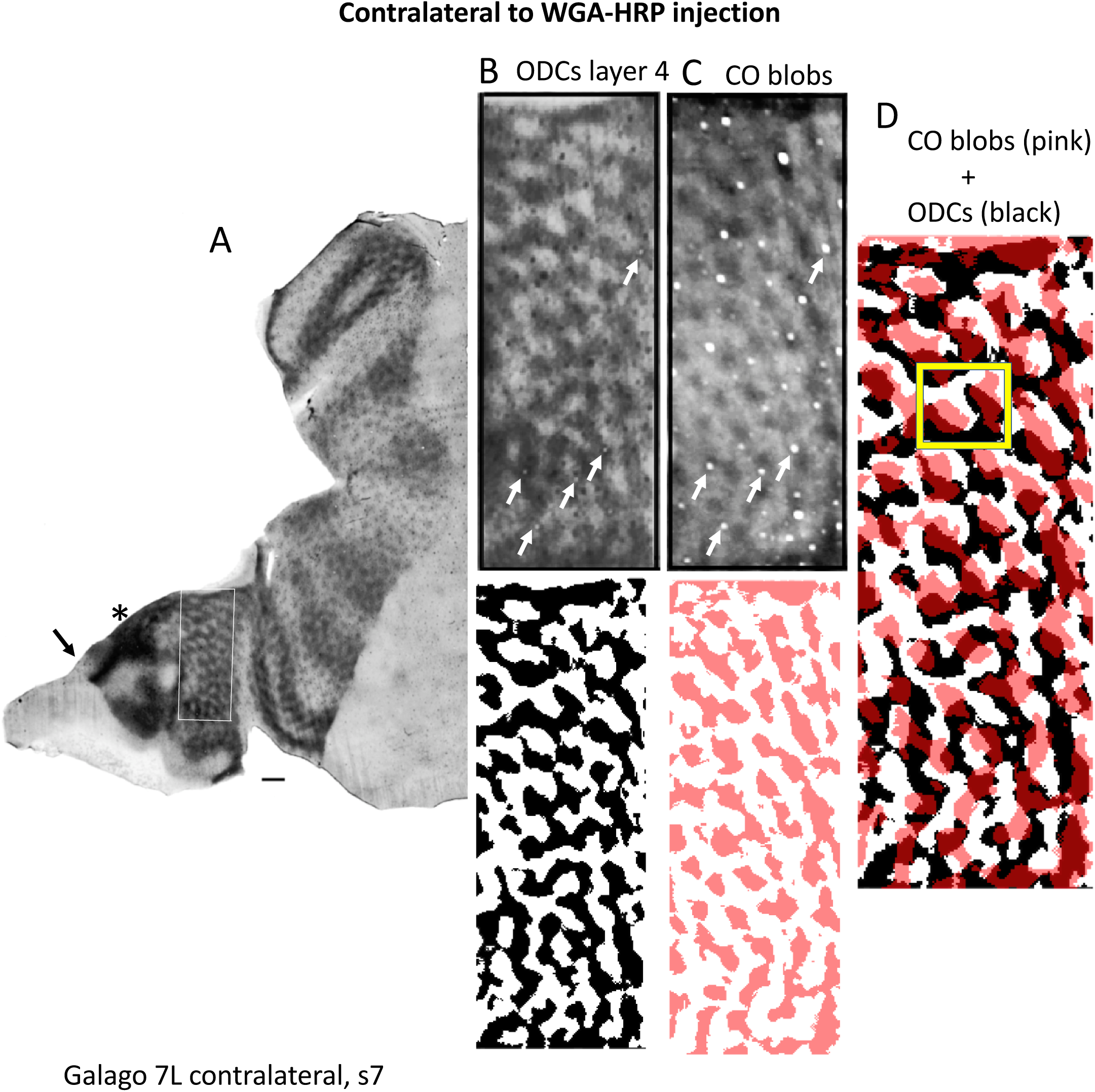
Correlation between the WGA-HRP labeling pattern in layer IV and the pattern of CO blobs in hemisphere contralateral to the eye injection in galago 7. A. Overall pattern of WGA-HRP labeling. The region outlined by the white box shows labeling in layer IV. This region is magnified in B. C. Pattern of CO blobs in the same region. White arrows in B,C indicate some of the blood vessels used to superimpose patterns in B and C. Corresponding thresholded patterns are shown below for ODCs in layer 4 (black) and CO blobs (pink). D. Superimposition of CO blobs (pink) over contralateral(black) and ipsilateral (white) ocular dominance columns in layer IV. The yellow box outlines a region used in Fig. 8. Scale bars in A = 1.0 mm.

**Figure 7.**
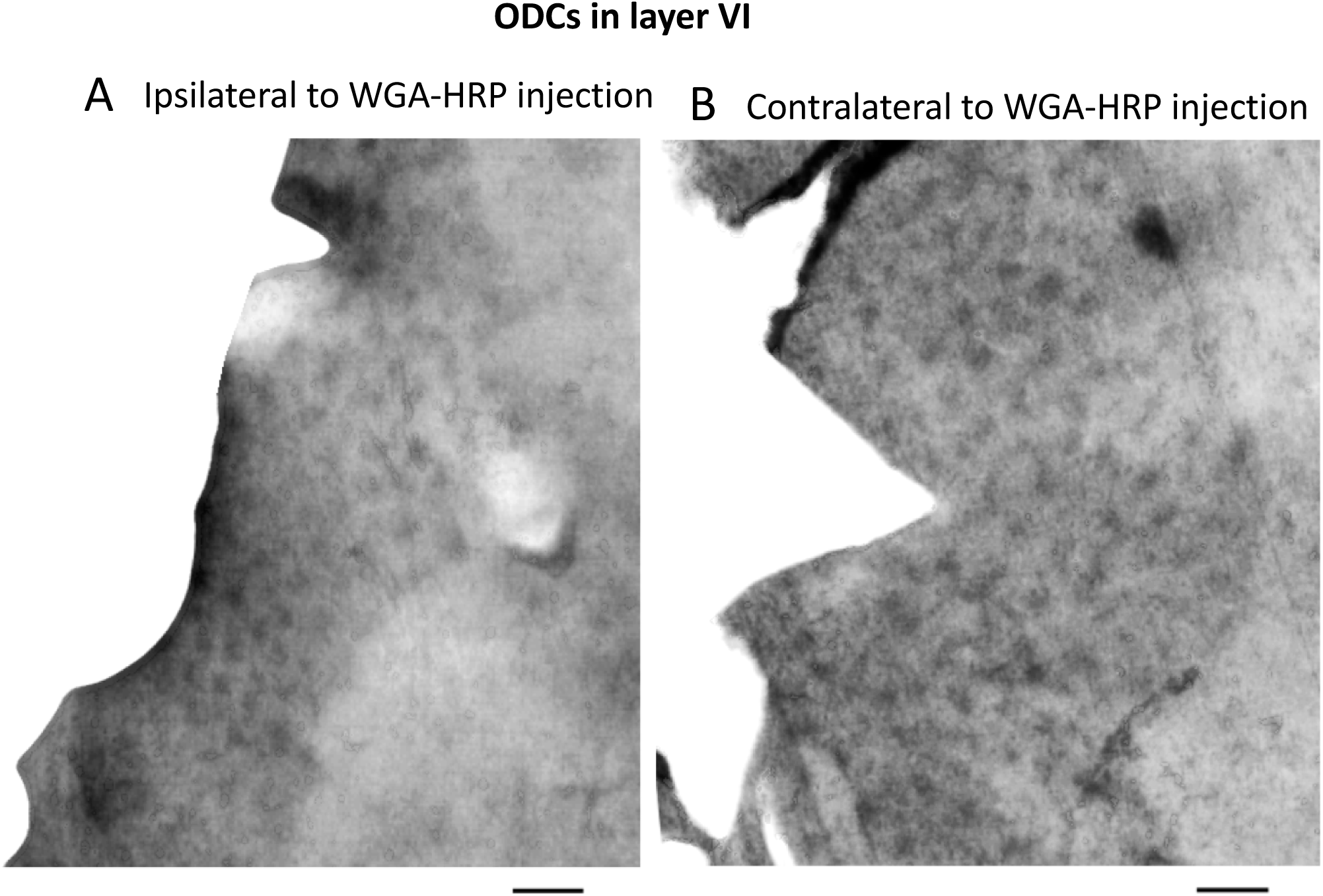
Overall pattern of WGA-HRP labeling in layer 6 in hemispheres ipsilateral (A) and contralateral (B) to the eye injection in galago 7. Scale bars = 1.0 mm.

**Figure 8.**
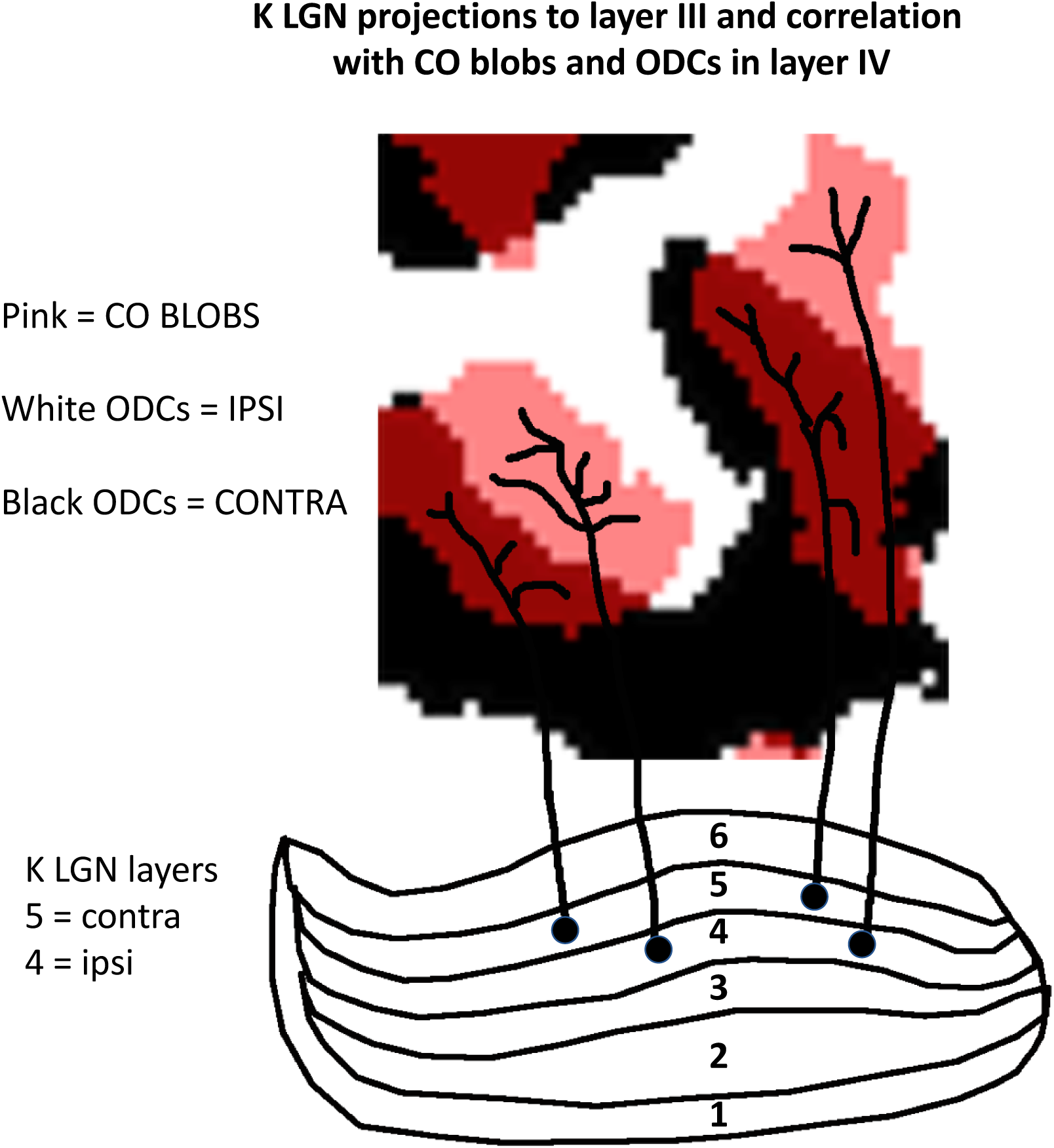
Diagram based on the case shown in Fig. 6D (region outlined in red in Fig. 6D). It illustrates that individual CO bobs (pink) overlap with labeled (black) and unlabeled (white) ODCs in layer 4. It also illustrates that the sizes of the arborizations of afferents from ipsilateral and contralateral K LGN layers (represented as branching black lines) may be constrained by the sizes of the CO blob subregions of the same eye dominance. Cells in ipsilateral and contralateral K LGN layers (layers 4 and 5, respectively) are represented as black dots.

We also examined 3 hemispheres from two additional cases injected with WGA-HRP into one eye (galago 3 and galago 6), and the results confirm our observations from the cases described above regarding the distribution of layer specific patterns of WGA-HRP labeling, and their correlation with CO blobs (data not shown).

## Discussion

### Overall pattern of layer IV ocular dominance columns in galago V1

We revealed, for the first time, that ODCs in galago layer 4 form a relatively continuous system of irregularly branching bands, about 200-250 um in width, in the hemispheres contralateral and ipsilateral to the injected eye. This pattern resembles the patterns of ODCs in cat layer IV (Anderson et al., 1988; Olavarria 2001), squirrel monkey (Horton and Hocking, 1996a) and owl monkey (Takahata et al., 2014). Moreover, as in these species, the bands in galago layer IV do not appear to have a bias in the distribution of orientations, and do not meet the border of V1 at right angles. In contrast, in macaque (e.g., Fig. 6 in Olavarria and Van Essen, 1997), and the New World monkey *Cebus apella* (Rosa et al., 1988), ODCs appear as largely parallel stripes that intersect the V1 border at right angles. One factor that may influence the orientation of ODCs is the emphasis on central visual representation in V1. Rosa et al., (1997) report that, in galago, the emphasis on central visual field representation in V1 is significantly less than in other primate species, but comparable to that of other non-primates, like cats. Another factor considered in previous studies is the overall shape of V1, and how the contralateral and ipsilateral visual fields are represented with respect to the main axes of V1. Anderson et al. (1988) point out that while the shape of V1 is elliptical in both macaque and cat, such that the ratio between long and short axes is nearly identical in both species, the horizontal meridian runs along the long axis of V1 in macaques, while in cats it runs along the short axis. This 90-degree rotation of the visual hemifield map in V1 would impose different constraints for accommodating the alternate mapping of contralateral and ipsilateral slices of the visual field, resulting in the pattern observed in the cat, in which the contralateral and ipsilateral strips alternate forming seemingly random patterns (Anderson et al., 1988, Olavarria, 2001).

Interestingly, in rats, as in cats, V1 is elliptical in shape and the horizontal meridian runs along the short axis of V1. Ocular dominance columns in this rodent form patches of various shapes and sizes that show no orientation bias with respect to the V1 border (Laing et al., 2015; Olavarria et al., 2021). In galago (Rosa et al., 1997) and squirrel monkey (Horton and Hocking, 1996a), V1 is nearly circular in shape, which could impose constraints leading to the development of irregularly branching eye specific bands. More recently, Nafajian et al. (2019) proposed a cortical model that generates variations in ocular dominance patterns from variations in local cortical retinotopy. They describe a common organizing principle across species that aligns the cortical axis of ocular dominance segregation with the axis of slowest retinotopic gradient, arguably accounting for ODCs patterns as diverse as those in macaques, squirrel monkey, cats and rats.

### CO blobs in galago V1 straddle the border of ocular dominance columns and receive segregated K LGN projections from both eyes

Lachica and Casagrande, (1992) injected the K LGN layers in galago with anterograde tracers and reported that terminations from individual K LGN axons form arbors of different sizes that are restricted to CO blobs, typically arborizing in only a fraction of the target CO blob. Our approach gave us the opportunity to examine the relationship between layer III patches and CO blobs in broad regions of both ipsilateral and contralateral hemispheres in the same animal. In each hemisphere, we found that virtually all CO blobs overlap with patches of K LGN projections in layer III. The overlap between layer III patches and CO blobs is partial, indicating that layer III patches occupy only a portion of the CO blob, in agreement with Lachica and Casagrande (1992). Our results imply that virtually all CO blobs receive K LGN input from both eyes. In addition, our data show that in some instances, the K LGN input extends into the neighboring interblob region. This observation would seem to be at odds with the report by Lachica and Casagrande (1992) that terminations of individual axons are restricted to CO blobs. We believe that the study by these authors does not necessarily contradict our results because we did observe many instances in which the layer III projections were largely or entirely restricted to the CO blobs. Moreover, Ding and Casagrande (1997) reported that, in owl monkeys, “the K terminals appear, in most cases, to extend slightly beyond the blob borders”. Lachica and Casagrande (1992) reconstructed a small sample of axons from parasagittal sections, and is possible that, by chance, the axon terminations they reconstructed were restricted to CO blobs. The function of K projections to CO blobs is discussed in Ding and Casagrande (1997), and Lachica et al. (1992), but at present, the possible function of K LGN projections to interblob regions in layer III remain to be investigated.

We also documented that CO blobs consistently straddle the border between neighboring ipsilateral and contralateral ODCs in layer IV, and our chi-square analysis showed that layer III patches are significantly correlated with layer IV ODCs of the same eye dominance. Together, our findings suggest that K LGN projections from each eye are segregated into ipsilateral and contralateral territories within the CO blobs. Support for this idea comes from previous physiological studies reporting that a high percentage of cells in CO blobs in galago V1 are monocular (DeBruyn et al., 1993). This idea is also consistent with our estimate that the average size of CO blobs is larger than the average size of layer III patches, while the average number of CO blobs per unit area is very similar to the average number of layer III patches labeled following an injection of WGA-HRP into one eye (Table 1). Figure 8, based on the case shown in Fig. 6D (region outlined in red in Fig. 6D), illustrates and summarizes these findings. It depicts that individual CO bobs (pink) overlap with labeled (black) and unlabeled (white) ODCs in layer 4. Figure 8 also proposes that the sizes of the arborizations of afferents from ipsilateral and contralateral K LGN layers (represented as branching black lines) may be constrained by the sizes of the CO blob subregions of the same eye dominance. Our finding that CO blobs consistently straddle the border between ODCs contrasts with the results of an optical imaging study of intrinsic signals in galago, which found no clear relationship between the distribution of CO blobs and the ocular dominance domains (Xu et al., 2005). The different outcomes may be attributed to differences in the methodology used in both studies.

In macaque monkeys (Horton, 1984; Blasdel and Salama, 1986; Tootell et al., 1988; Bartfeld and Grinvald, 1992; Yoshioka et al., 1996) and humans (Adams et al., 2007), CO blobs are centered on ocular dominance columns and appear to be dominated by the monocular input represented in the underlying ODC (Livingstone and Hubel, 1984). The model we propose in galago (Fig. 8) can be hypothetically transformed into a model isomorphic with the macaque model by dividing the galago CO blobs into their two eye dominance regions, and then displacing each region toward the center of the underlying ODC of the same eye dominance. It would be interesting to determine whether there is a phylogenetic relation between the models for galago and macaque monkey. However, the fact that a clear and consistent correlation between CO blobs and ODCs has not been found in New World primates (Roe et al., 2005; Kaskan et al., 2007; Takahata et al., 2014) suggests that other structural and functional relationships between CO blobs and ODCs may operate in these primates. In owl monkey, Takahata et al., (2014) found that ODCs adopt the form of interconnected bands or stripes, whereas they tend to be patchy in peripheral regions of V1. Moreover, these authors found that CO blobs straddle the border of ODCs in central regions of V1, but they are centered on ODCs patches in peripheral V1. The possibility that differences in the balance of left and right eye dominance in central vs. peripheral regions of V1 influence the correlation between CO blobs and ODCs is discussed in Takahata et al. (2014).

### Comparison with cats

Previous studies have shown that parallel visual processing streams (Leventhal et al., 1985; reviewed in Casagrande and Norton, 1991) and CO blobs in V1 (Boyd and Matsubara, 1996; Murphy et al., 1995) exist in cats. As in primates, parallel retino-cortical pathways in cats originate from three distinct classes retinal ganglion cells, named X, Y and W, which, on the basis of their appearance and physiological properties, have been compared to P, M and K ganglion cells in primates, respectively (Leventhal et al., 1985; reviewed in Casagrande and Norton, 1991). Boyd and Matsubara (1996) report that projections from W cells (K-like) in cat LGN layers C1 and C2 are patchy in layer III of V1, and that the patches align with CO blobs, resembling the projection from K LGN layers to CO blobs described in galago (Lachica and Casagrande, 1992; Casagrande and Kaas, 1994; present study). It remains to be investigated whether all CO blobs in cat V1 receive W LGN projections from both eyes, as we found in this study to be the case for K LGN projections in galago. Regarding the position of CO blobs with respect to ODCs in cat layer IV in V1, Murphy et al. (1995) reported that the location of some CO blobs in V1 is not above the center of labeled ODCs. However, after considering that, relative to the width of ODCs, the diameter of CO blobs in cats is greater than in macaques, and that in cats there is some overlap between ipsilateral and contralateral ODCs, these authors proposed that the functional organization of the blobs is similar in cats and macaque monkeys. Galagos also resemble cats in the distribution of callosal connection in V1. In galagos (Weyand and Swadlow, 1980; Cusick et al., 1984; Beck and Kaas, 1994, as in cats (Boyd and Matsubara, 1994; Olavarria, 2001), callosal connections in V1 are robust and occupy a broad area extending medially from the V1/V2 border. In cats, callosal cells correlate with contralateral ODCs within the narrow V1/V2 transition zone (TZ), and with ipsilateral ODCs in regions of areas V1 and V2 located outside the TZ. Finding that ODCs in galago show the same association with visual callosal connections would suggest that the retinally driven mechanism that is thought to specify the distribution of callosal connections in cat V1 (Olavarria, 2001) also operates in galago.

In conclusion, our study is significant for mainly two reasons. It incorporates the prosimian galago into the group of primate and non-primate species that have well-defined pattern of ODCs in layer IV of V1. In addition, our results suggest that ocular dominance is a key factor orchestrating the close relationship that we observed between ODCs in layer IV, CO blobs, and K LGN projections to CO blobs from both eyes. Our findings in galago should facilitate, and serve as a reference for, future studies of the development, plasticity and function of ocular dominance domains and parallel visual pathways in prosimian primates.

## Conflict of Interest

Authors declare no conflict of interest

## Acknowledgments

We greatly acknowledge the guidance and help by Dr. Casagrande, and the technical support by personnel in the labs of Dr. Casagrande and Dr. Kaas. This research was supported by National Institutes of Health grant numbers: EY01778, EY02686, EY09343 (to JHK), EY016045, NS070022 (to JFO), and by National Natural Science Foundation of P. R. China 32170992, 31872767 (to TT).

## Notes

### Competing Interest Statement

The authors have declared no competing interest.

